# Amino acid and nucleotide metabolism shape the selection of trophic levels in animals

**DOI:** 10.1101/2021.07.01.450789

**Authors:** Rosemary Yu, Hao Wang, Jens Nielsen

## Abstract

What an animal eats determines its trophic level (TL) in the food web. The diet of high-TL animals is thought to contain more energy because it contains higher levels of lipids. This however has not been systematically examined in the context of comprehensive metabolic networks of different animals. Here, we reconstruct species-specific genome-scale metabolic models (GEMs) of 32 animals, and calculate the maximum ATP production per unit of food for each animal. Surprisingly, we find that ATP production is closely associated with metabolic flux through central carbon metabolism and amino acid metabolism, while correlation with lipid metabolism is low. Further, metabolism of specific amino acids and nucleotides underlie maximum ATP production from food. Our analyses indicate that amino acid and nucleotide metabolism, rather than lipid metabolism, are major contributors to the selection of animal trophic levels, demonstrating that species-level metabolic flux plays key roles in trophic interactions and evolution.

## Introduction

The choice of food used by an organism for nutritional intake, and the breakdown of nutrients to generate energy in the form of adenosine triphosphate (ATP), are fundamental to living systems. Animals exhibit a wide diversity of dietary choices which place them at different trophic levels (TL) in a food chain or food web (Fig 1A). In terms of food availability, plants and algae/phytoplankton are exceptionally abundant and stable dietary resources, and have been so across evolutionary history ^1–3^. Moreover, a plant-based diet is estimated to be >10-fold more ecologically efficient than an animal-based diet, since only a fraction of the energy available in the biomass of a given TL is transferred to the next upper TL ^4–7^. Consistently, the evolutionary diversification rate of herbivores has been found to be faster than that of carnivores ^8^, while specialization in carnivory appears to be unstable, as it is associated with short extinction times and repeated ecological replacements ^9,10^. Surprisingly, however, recent large-scale studies have shown that a relatively small proportion of animal species are herbivores (32-43%), whereas 57-66% of species have a diet consisting, partially or completely, of other animals ^8,11^. This suggests a selective advantage in an animal-based diet, which thereby favors the selection of high-TL animals.

**Figure 1.**
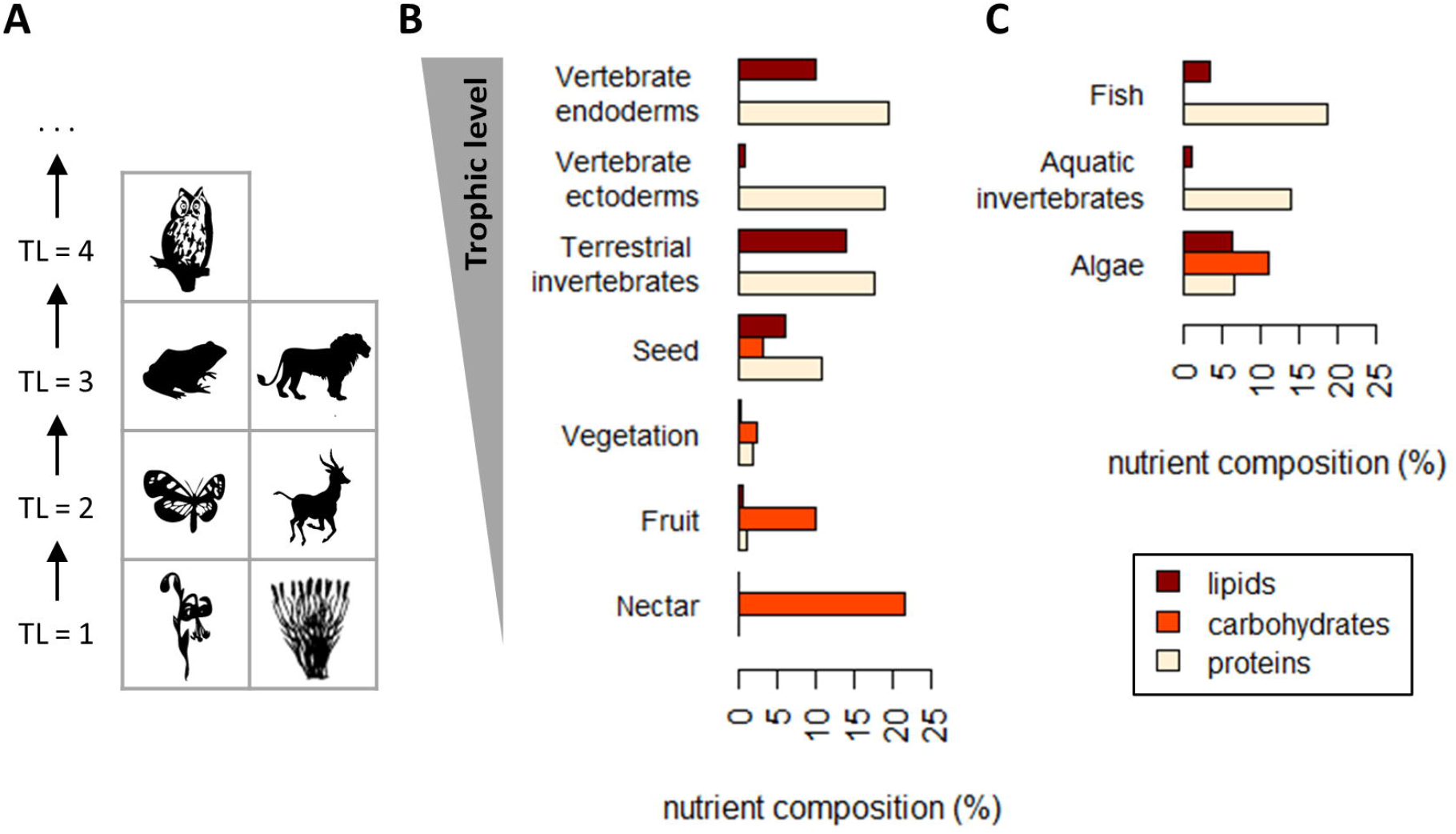
Animal trophic levels (TL) and nutrient composition of different diet types. **A**, two examples of food chains are shown to demonstrate the concept of TL. **B-C**, the composition of the three major dietary macronutrients (lipids, carbohydrates, and proteins) in different diet types of terrestrial (**B**) and aquatic (**C**) species, order by trophic level, is given in % (g/g wet weight). Diet types are as collated in EltonTraits (Wilman *et al*, 2014).

The traditional assumption (e.g. ref ^12^) for this selection pressure is that an animal-based diet contains more lipids, whereas a plant-based diet contains less lipids but more carbohydrates (Fig 1B-C). As the energy density of lipids is higher than carbohydrates ^13,14^, this suggests that high-TL animals would be able to extract more energy per unit of food, consistent with the observation that carnivores spend less time feeding than similar-sized herbivores ^15^. In other words, dietary lipid to carbohydrate ratio is thought to be the main determinant of the amount of energy obtained by an organism per unit of food. However, in living systems energy is extracted as ATP through a complex network of metabolic reactions, and the bottleneck(s) of ATP production in the context of the metabolic network of different animals is unknown. Moreover, there is substantial differences in dietary protein content with respect to TL (Fig 1B-C), which has an energy density comparable to that of carbohydrates, which can further contribute to ATP production. A closer examination of the metabolic determinants in trophic selection is therefore warranted.

A genome-scale metabolic model (GEM) is a constraint-based modeling framework wherein the metabolic network of an organism is represented mathematically ^16,17^. Simulations using GEMs are based on the concept of flux balance analysis (FBA) ^16^, and can be used to calculate the production level of a metabolite with given constraints in the intake of nutrients. GEMs for microorganisms have been used for such simulations for many years ^18,19^, and recently a unified Human-GEM, containing 8,378 metabolites and 13,072 reactions, has also been published ^20^, following >15 years of a concerted community effort.

Here, by using Human-GEM as a template ^20^, we report the reconstruction of GEMs for 32 animals and their use for performing FBA simulations at a range of TLs. Imposing constraints on these GEMs based on the dietary composition of carbohydrates, lipids, and proteins allows the simulation of ATP production per unit of food in each species. Interestingly, simulation results show that ATP production in relation to TL is associated with dietary protein content, rather than dietary lipid content. We further show that TL is substantially correlated with metabolic flux through most reactions in central carbon metabolism and amino acid metabolism (in particular Lys, Trp, Tyr, Ile, Leu, and Val), but only a small proportion of reactions in lipid metabolism. Finally, we find that metabolic pathways of specific amino acids (His metabolism, Thr-Gly conversion, and Asn-Asp conversion) and nucleotides underlie maximum ATP production from food. Taken together, our analyses indicate that amino acid and nucleotide metabolism play major roles in shaping the metabolism of high-TL animals, and suggest that species-level metabolic flux can play key roles in trophic interactions and evolution.

## Results

### Dietary nutrient composition

We reconstructed species-specific animal GEMs for a total of 32 species, including 22 terrestrial and 11 aquatic organisms, with TL ranging from 2 to 4.2 (Supplementary Table 1; see Methods section for data source and related calculations). To constrain the models based on dietary nutrient composition, we calculated the % (g/g wet weight) of dietary carbohydrates, lipids, and proteins for each species (Supplementary Table 1), based on attributes mined from EltonTraits ^21^ and FishBase ^22^ (see Methods section). Consistent with previous literature, dietary carbohydrate content decreases with TL (Fig 2A). At around TL = 2, the dietary carbohydrate content varies between 2-8%, as there are large differences in the carbohydrate content in different parts of plants (Fig 1B-C). The diet of the herbivorous (TL = 2.05) fish *Oreochromis niloticus* (Nile tilapia) contains 11% carbohydrates, reflecting the high carbohydrate content in algae ^23^. Above around TL = 3, dietary carbohydrates of both terrestrial and aquatic species decreased below 1% (Fig 2A). In contrast, the dietary content of proteins increases with TL, from 2% to 19%, and data from both terrestrial and aquatic species follow closely the same trajectory (Fig 2B). The dietary content of lipids also increases with TL, however data from terrestrial and aquatic species clearly follow distinct trajectories, with terrestrial species reaching up to 10% dietary lipids at high TL, while aquatic species reaching only 4% (Fig 2C). The diet of *Oreochromis niloticus* contains 6% lipids, again reflecting the biomass composition of algae ^23^, which places the dietary lipid content of this fish in the trajectory of terrestrial animals (Fig 2C).

**Figure 2.**
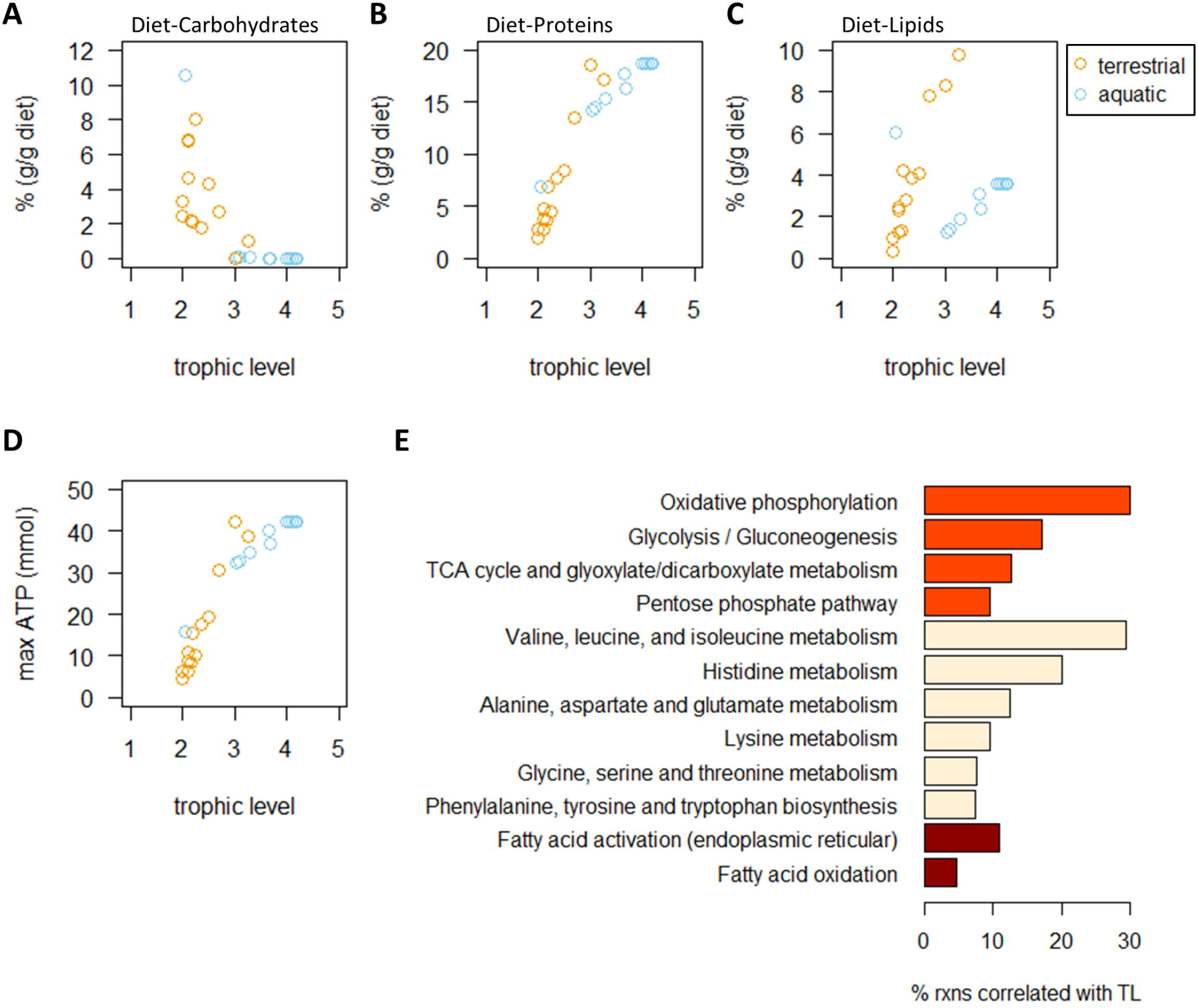
Dietary nutrient composition and maximum ATP production. **A-C**, dietary carbohydrates (**A**), proteins (**B**), and lipids (**C**) of 33 species (32 animals plus human), is given with respect to the species-specific TL. Terrestrial species are in orange and aquatic species are in blue. **D**, the maximum ATP production as simulated using species-specific GEMs and constrained to reflect 100 g of food intake. **E**, the metabolic subsystems wherein a high % of reactions show high correlation with TL in FBA and random sampling analyses. Subsystems related to carbohydrate metabolism, amino acid metabolism, and fatty acid metabolism are shown.

### ATP production is not associated with dietary lipid content

We used FBA to simulate the maximum ATP production for each species, given the constraints in dietary carbohydrates, proteins, and lipids. The reactions of O_2_ uptake, CO_2_ production, as well as exchange reactions of H_2_O, protons, and metal ions, were unconstrained. For nitrogenous waste, the exchange reactions of urea, urate, allantoin, or NH_3_, were either unconstrained or constrained to 0, to reflect the species-specific waste product (see Methods section). All other exchange reactions were constrained to 0. We found that ATP production increased with increasing TL (Fig 2D), and importantly, it is not grouped into distinct trajectories depending on the habitat of the animals, as is the case with dietary lipid content (Fig 2C). This indicates that, contrary to the traditional assumption ^12^, lipid content is not the limiting dietary nutrient in ATP production in relation to TL.

To further examine the metabolic constraints of ATP production from the diet in different animals, we constrained the ATP production reaction in each GEM to the maximum calculated value (Fig 2D), and implemented FBA with random sampling ^24^ to obtain a set of 1,000 possible flux distributions within the feasible region. The average flux of each reaction (Supplementary Table 2) was then correlated with TL, and hereby we found that a large proportion of reactions in metabolic subsystems related to central carbon metabolism and amino acid metabolism are highly correlated with TL (Fig 2E). In contrast, the pathways related to lipid metabolism have relatively few reactions correlating with TL (Fig 2E), in agreement with the result that ATP production is not constrained by the dietary lipid content of animals at different TLs.

### Metabolic fluxes correlated with TL

We then examined the specific reactions and pathways that are found to be highly correlated with TL. In the glycolysis-gluconeogenesis pathway, almost all reactions in ‘lower’ glycolysis, involving a chain of metabolic conversions between 3-carbon molecules, are highly correlated with TL (Fig 3A); whereas reactions in ‘upper’ glycolysis, involving the metabolism of 6-carbon molecules, are not. The reaction that connects upper and lower glycolysis (HMR_4375r), which converts the 6-carbon molecule fructose 1,6-bisphosphate (F1,6-bP) to 3-carbon molecules dehydroxyacetone phosphate (DHAP) and glyceraldehyde 3-phosphate (G3P), shows borderline correlation with TL with *ρ*_Spearman_ = 0.75 (Fig 3A). In particular, lower glycolysis generally carries positive flux in low-TL species, consistent with the use of glycolysis to metabolize the high levels of carbohydrates in the diet of these organisms (Fig 2A). In high-TL species, this pathway carries negative flux (Fig 3A), indicating the use of gluconeogenesis to synthesize other metabolic intermediates, at a cost of ATP.

**Figure 3.**
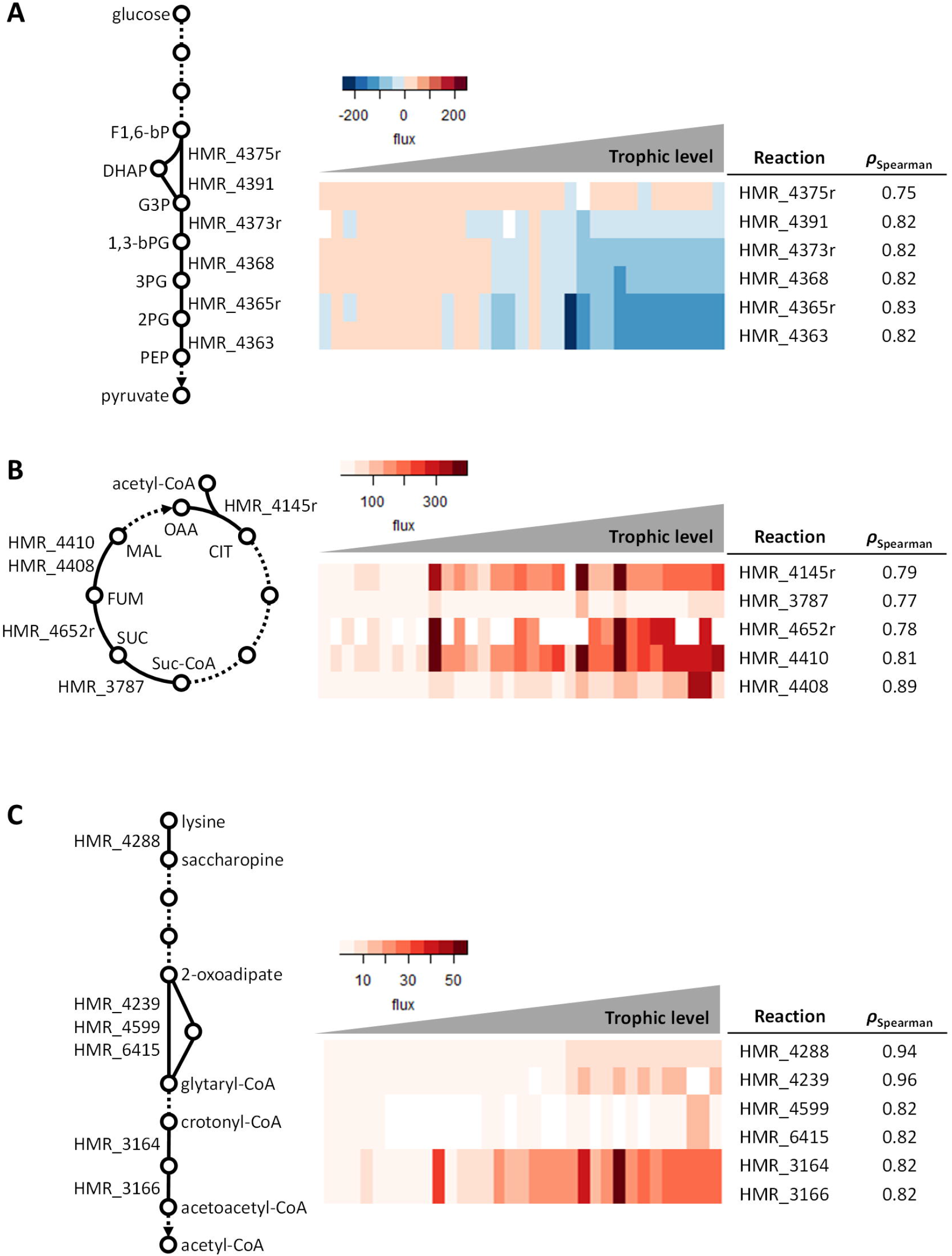
Select metabolic pathways with a high % of reactions showing high correlation with TL. The pathways of glycolysis (**A**), TCA cycle (**B**), and lysine degradation to acetyl-CoA (**C**) are shown. Reactions that have high correlation with TL are represented by solid lines, and the GEM reaction ID is given. Reactions that have low correlation with TL are represented by dotted lines. Key metabolites in each pathway are indicated. F1,6-bP, fructose 1,6-bisphosphate. DHAP, dihydroxyacetone phosphate. G3P, glyceraldehyde 3-phosphate. 1,3-bPG, 1,3-bisphosphoglycerate. 3PG, 3-Phosphoglycerate. 2PG, 2-Phosphoglycerate. PEP, phosphoenolpyruvate. OAA, oxaloacetate. CIT, citrate. Suc-CoA, succinyl-CoA. SUC, succinate. FUM, fumarate. MAL, malate.

In the tricarboxylic acid (TCA) cycle, we found that the first step of the pathway (HMR_4145r) catalyzing the entry of acetyl-CoA into the cycle, as well as several steps in the second half of the cycle converting succinyl-CoA (Suc-CoA) to malate (MAL), are highly correlated with TL (Fig 3B). This is likely related to high levels of acetyl-CoA and Suc-CoA entering the TCA cycle in high-TL species. As the high dietary protein levels (Fig 2B) provides disproportionate levels of amino acids, many of the amino acids are deaminated and converted into acetyl-CoA or succinyl-CoA, entering the TCA cycle to generate ATP. As an example, Fig 3C shows the degradation of lysine to acetyl-CoA, wherein several steps are highly correlated with TL. The degradation of aromatic amino acids to acetyl-CoA, and branched chain amino acids to succinyl-CoA, all follow similar trends (Supplementary Table 2).

### ATP production is constrained by nucleotide metabolism

In amino acid metabolic pathways, beyond the increase in the degradation of amino acids to acetyl-CoA and Suc-CoA within increasing TL (Fig 3B-C and Supplementary Table 2), we also found that TL is highly correlated with several amino acids being shunted towards the synthesis of metabolic intermediates in nucleotide synthesis. In particular, nearly all steps in the conversion of histidine to 10-formyl THF, which enters the nucleotide metabolic pathway at two distinct steps, are highly correlated with TL, with an overall *ρ*_Spearman_ = 0.96 (Fig 4A). The conversion of threonine to glycine, and asparagine to aspartate, are similarly highly correlated with TL (*ρ*_Spearman_ = 0.94 and 0.85 respectively), consistent with an increased supply of GAR (Glycineamideribotide) and SAICAR (Phosphoribosylaminoimidazolesuccinocarboxamide) to the nucleotide metabolic pathway (Fig 4A). Moreover, reactions in the entire pathway of IMP (inosine monophosphate) production from PRPP (phosphoribosyl pyrophosphate), all exhibit high correlations with TL (*ρ*_Spearman_ = 0.96), suggesting that nucleotide metabolism, rather than the lipid metabolism, plays a significant role in ATP production in animals at different TLs.

**Figure 4.**
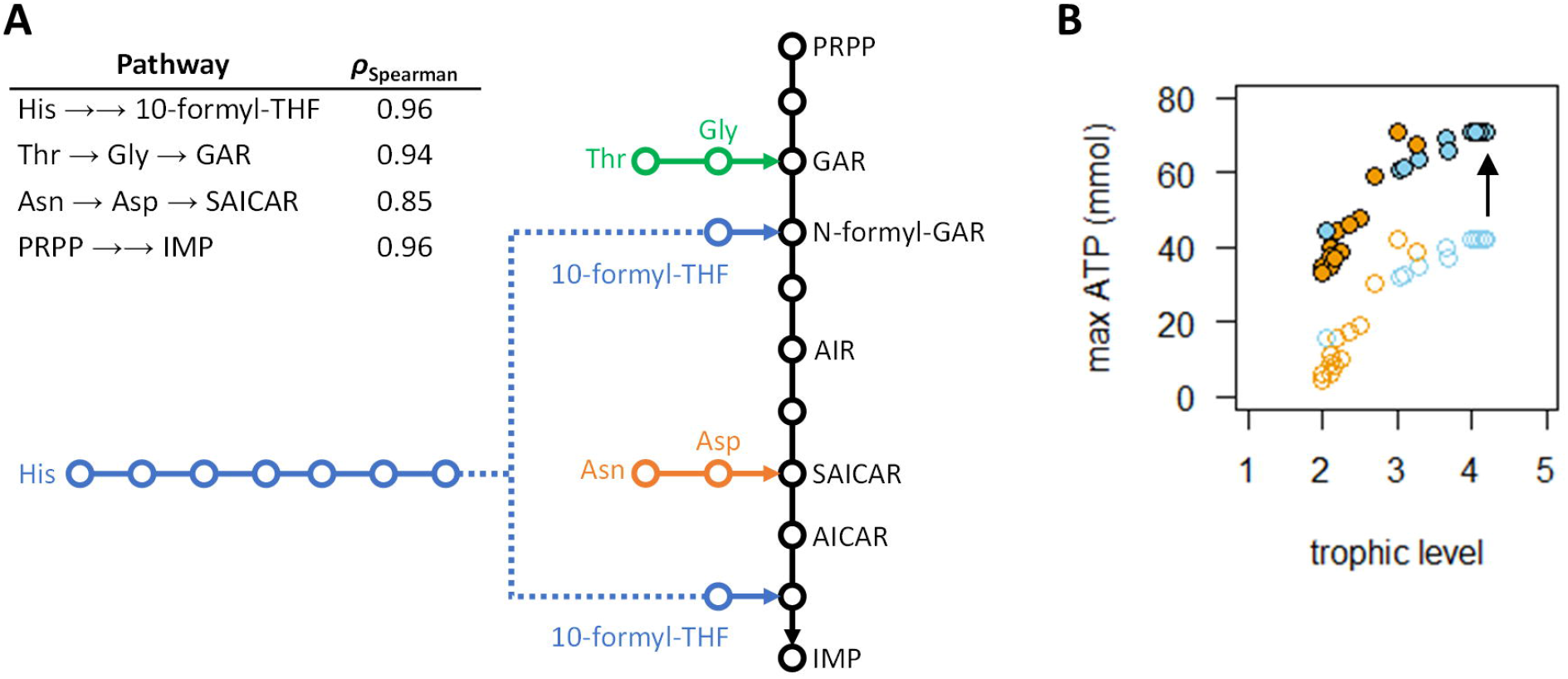
ATP production is constrained by nucleotide metabolism. **A**, the nucleotide synthesis pathway from several precursors is shown. Reactions that have high correlation with TL are represented by solid lines. Key metabolites are indicated. Colors separate the main pathway (black) from the synthesis of different precursors. PRPP, phosphoribosyl pyrophosphate. GAR, 5’-phosphoribosylglycinamide. N-formyl-GAR, 5’-phosphoribosyl-N-formylglycinamide. AIR, 5’-phosphoribosyl-5-aminoimidazole. SAICAR, 1-(5’-phosphoribosyl)-5-amino-4-(N-succinocarboxamide)-imidazole. AICAR, 1-(5’-phosphoribosyl)-5-amino-4-(N-succinocarboxamide)-imidazole. IMP, inosine monophosphate. 10-formyl-THF, 10-formyltetrahydrofolate. **B**, the maximum ATP production as simulated using species-specific GEMs and constrained to reflect 100 g of food intake, with 10% of the food allowed to be adenosine monophosphate (AMP).

While it makes intuitive sense that ATP production is constrained by the metabolic pathway catalyzing the synthesis of nucleotides, these results do not account for any nucleotides that are already available as a part of the diet (free or bound in DNA/RNA). As measurements of nucleotide content in different dietary sources are scarce, we addressed this by constraining our model to allow a dietary AMP intake of up to 10% (g/g wet weight). We then used GEMs to simulate the maximum ATP production in each species as before. Our results show that allowing for up to 10% of dietary AMP led to an up-shift in the maximum ATP production in each species by ^~^29 mmol, but did not alter the overall trajectory of ATP production with respect to TL (Fig 4B). Data from terrestrial and aquatic species remained on the same trajectory, instead of separating into distinct groups which would be the case if dietary lipids were the constraining nutrient for ATP production (Fig 2C). We therefore conclude that amino acid and nucleotide metabolism play key roles in the production of ATP from food, which in turn contributes to the evolutionary selection of animals at high TL.

## Discussion

We reconstructed species-specific GEMs for 32 different animals and simulated the maximum ATP production per unit of food. Our analyses show that dietary protein content, rather than dietary lipid content, supports an increased ATP production that correlates with increasing TL. Results further indicate that ATP production from food is limited by the metabolic pathways of histidine metabolism, threonine-glycine conversion, asparagine-aspartate conversion, and nucleotide metabolism. We therefore conclude that amino acid and nucleotide metabolism are major contributors to the evolutionary selection of animal-based diets and high-TL animals, contrary to the traditional assumption ^12^ of lipid content being the primary selection pressure.

Despite the prevalent use of dietary lipid content to explain the selection of high-TL animals, this does not always fit existing data. For instance, while animal-based diets are generally thought to contain higher levels of both proteins and lipids compared to plant-based diets, this is only true of terrestrial environments. In aquatic environments, the lipid content of fish is known to be very low, whereas the biomass composition of algae contains higher levels of lipids ^23^ than fish ^22,25^ or aquatic invertebrates ^26^. As such, in aquatic environments, the diet of herbivorous species actually contains more lipids than omnivorous or carnivorous species. Therefore, dietary lipid content cannot explain the selection of high-TL species in aquatic ecosystems, suggesting that alternative factors are involved. Of the three main energy-carrying macronutrients in the diet, only the protein content is higher in omnivorous/carnivorous fish than in herbivorous fish (here represented by *Oreochromis niloticus*, Fig 2A-C), suggesting that dietary protein content plays a key role in the selection of high-TL species, in line with our GEM simulation results.

Our results also show that, when a dietary nucleotide content of 10% is included as a simulation constraint upper bound, this leads to an up-shift in the maximum ATP production in all species, by an equal amount of ^~^29 mmol (Fig 4B). In reality, however, this up-shift is likely not constant across all species, but rather increases with increasing TL. This is because nucleotide levels generally track with protein levels, in part because the majority of RNA in living cells is ribosomal RNA which is closely associated with ribosomal proteins ^27^. Thus, diets of high-TL animals would contain higher levels of both proteins and nucleotides, leading to a steeper slope of ATP production with respect to TL. Moreover, in low-TL animals, plant-based diets could contain toxins or anti-nutrients which limit the bioavailability of nutrients ^28,29^. For example, trypsin inhibitors and hemagglutinins found in legumes can reduce the digestibility of proteins and amino acids by up to 50%; tannins found in cereals have similar effects by up to 23%; phytates in oilseeds, by up to 10%; and many more ^29^. These factors would further increase the steepness of the slope of ATP production with respect to TL, providing additional selection pressure for high-TL animals.

In addition to amino acid and nucleotide metabolism, our analyses show that metabolic flux through lower glycolysis and the second half of the TCA cycle, are also correlated with TL. Of particular note is that with increasing TL there is a reversal of the direction of flux in lower glycolysis, catalyzing glycolysis in low-TL animals, and gluconeogenesis in high-TL animals (Fig 3A). However, enzymes in this pathway are highly conserved across all organisms ^30,31^, suggesting that the versatile use of this pathway to metabolize dietary nutrients in both directions is independent of enzyme properties and likely reflects the biochemistry of the pathway itself. Indeed, recently it has been shown that, out of hundreds of (theoretically) feasible alternative biochemical paths connecting G3P to pyruvate, the extant lower glycolysis is the optimal solution which carries the maximum flux for both glycolysis and gluconeogenesis under biologically relevant conditions ^32^. For all other pathways found to correlate with TL, in particular the amino acid and nucleotide metabolism pathways involved in maximizing ATP production, whether adaptations in enzyme properties or pathway optimality underlies the selection of animal trophic levels remains an interesting open question.

## Methods

### GEM reconstruction

The animal GEMs were generated via an orthology-based approach, by using the RAVEN 2.0 package ^33^ and the Human-GEM version 1.7 ^20^ as a template. The annotated orthologs and paralogs associated from human to other animal species were retrieved from the Ensembl database version 103 ^34^.

### Diet type and TL calculations

Diets of terrestrial animals and whales (*Delphinapterus leucas* and *Physeter catodon;* see Supplementary Table 1) are obtained from EltonTraits ^21^ which contains the percent usage of 10 diet types. The diet type “Inv” (invertebrates) is further split to differentiate the usage of aquatic invertebrates or terrestrial invertebrates, based on “Food Habits” data mined from Animal Diversity Web ^35^, to a total of 11 diet types. Diet types are mined at the genus level, to account for missing data in Animal Diversity Web. For genus with food habits of insects, terrestrial non-insect anthropoids, or terrestrial worms, terrestrial invertebrate usage is equal to Diet-Inv. For genus with food habits of aquatic or marine worms, aquatic crustaceans, echinoderms Cnidarians, other marine invertebrates, or zooplankton, aquatic invertebrate usage is equal to Diet-Inv. For the food habit of mollusks, aquatic invertebrate usage is equal to Diet-Inv only for genus with food mollusks of both mollusks and fish, in order to exclude species that eat snails. If a genus is found to eat both aquatic and terrestrial invertebrates, usage of Diet-Inv is split in half into terrestrial and aquatic invertebrate usage. If no data is available, Diet-Inv is assumed to be terrestrial invertebrate usage. The *TL_j_* of each diet type *j* is then taken as follows: fruit, nectar, seed, and plant, *TL_j_* = 1; terrestrial invertebrates, *TL_j_* = 2; vertebrate endoderms, vertebrate ectoderms, and fish, *TL_j_* = 2.5. The *TL_i_* of each species *i* is then calculated as follows ^36^:

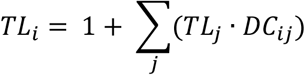

where *DC_ij_* represents the fraction of *j* in the diet of *i* ^21^.

For aquatic species except whales (*Delphinapterus leucas* and *Physeter catodon), TL_i_* is obtained from FishBase ^22^. The *TL_j_* of each diet type *j* is taken as follows: algae, *TL_j_* = 1; aquatic invertebrates, *TL_j_* = 2; fish, *TL_j_* = 3 for species at *TL_i_* between 3 and 4, and *TL_j_* = 4 for species at *TL_i_* between 4 and 5. The fraction of *j* in the diet of *i* (*DC_ij_*) is then calculated as above ^36^.

### Dietary nutrient composition

The dietary composition of carbohydrates, lipids, and proteins for the 11 diet types (see diet type and TL calculations section) are mined from existing knowledge bases as follows: fruit ^26^; nectar ^37,38^; seed, ^26^; plant, ^26^; algae, ^23^; aquatic invertebrates, ^26^; terrestrial invertebrates, ^39,40^; vertebrate endoderms, ^21,26^; vertebrate ectoderm, ^26,41^; vertebrate fish, ^22,25^. Vertebrate general/unknown and scavenge are taken as the average of vertebrate endoderm, ectoderm, and fish.

### FBA and random sampling

Constraints of each GEM are imposed as follows: for dietary carbohydrates, lipids, and proteins (CLP), both the upper bound (ub) and the lower bound (lb) of the glucose uptake reaction, lipid pool uptake reaction, and protein pool uptake reaction were constrained to the % (g/g) CLP in the diet (Supplementary Table 1), after conversion to mmol by the previously estimated molecular weight of the pool metabolites ^20^. For O_2_ uptake, the ub is constrained to 0, and the lb is constrained to -Inf (negative infinity). For CO_2_ production, the ub is constrained to Inf, and the lb is constrained to 0. Nitrogenous waste production are as follows: Simian primates (including humans) ^42^ excrete urate and urea; non-Simian mammals excrete urea and allantoin; and fish excrete urea and NH_3_. For each species, then, the allowed nitrogenous waste is constrained with ub to Inf and lb to 0. For water, proton, and metal ions (zinc, selenate, sulfate, sodium, magnesium, lithium, potassium, iodide, ferrous ion, ferric ion, cupric ion, phosphate, chloride, and calcium), ub is constrained to Inf and lb is constrained to -Inf. ATP production is constrained with lb to 0 and ub to Inf. All other exchange reactions are constrained to 0.

MATLAB R2019b (MathWorks, Inc., Natick, MA) with Gurobi solver (Gurobi Optimizer, Beaverton, OR) in the COBRA toolbox ^43^ was used for all GEM simulations. In FBA, ATP production is set to be the objective function, and Inf (or -Inf) are converted to 1000 (or −1000) to avoid loops. In FBA with random sampling, ATP production is constrained with both lb and ub equal to the maximum ATP production calculated in the first iteration of FBA, followed by 1000 random samplings of a pair of randomly weighted objective functions, as implemented in the RAVEN toolbox ^33^. Spearman correlation (*ρ*_Spearman_) between the mean of the 1000 random sampling results for each reaction in each species, and the TL of the species, is then calculated. Reaction flux is considered highly correlated with TL if *ρ*_Spearman_ > 0.75 or *ρ*_Spearman_ < −0.75, and if the number of species where the reaction carries non-0 flux is > 17 (i.e. 50% of all species considered). In subsystem analysis, subsystems wherein > 2 reactions are highly correlated with TL are considered.

## Data availability

Diet and TL data on the species studied are in Supplementary Table 1. Processed simulation results are in Supplementary Table 2. Species-specific GEMs and related data are available in the GitHub repository at https://github.com/SysBioChalmers/GEMsforTrophicLevels.

## Code availability

Custom scripts are available in the GitHub repository at https://github.com/SysBioChalmers/GEMsforTrophicLevels.

## Acknowledgements

This research was supported by funding from the Novo Nordisk Foundation (grant number NNF10CC1016517) and the Knut and Alice Wallenberg Foundation. Open access funding is provided by Chalmers University of Technology.

## Author contributions

R.Y., H.W., and J.N. conceived the study. R.Y. and H.W. designed and performed the analyses. J.N. supervised the study. All authors wrote the manuscript.

## Competing interests

The authors declare no competing interests related to this work.

## Materials and Correspondence

Correspondence to Jens Nielsen

## Supplementary information

Supplementary Table 1-2.

## Notes

https://github.com/SysBioChalmers/GEMsforTrophicLevels

### Competing Interest Statement

The authors have declared no competing interest.

